# Dynamics of natural selection preceding human viral epidemics and pandemics

**DOI:** 10.1101/2025.02.26.640439

**Authors:** Jennifer L. Havens, Sergei L. Kosakovsky Pond, Jordan D. Zehr, Jonathan E. Pekar, Edyth Parker, Michael Worobey, Kristian G. Andersen, Joel O. Wertheim

## Abstract

Using a phylogenetic framework to characterize natural selection, we investigate the hypothesis that zoonotic viruses require adaptation prior to zoonosis to sustain human-to-human transmission. Examining the zoonotic emergence of Ebola virus, Marburg virus, influenza A virus, SARS-CoV, and SARS-CoV-2, we find no evidence of a change in the intensity of natural selection immediately prior to a host switch, compared with typical selection within reservoir hosts. We conclude that extensive pre-zoonotic adaptation is not necessary for human-to-human transmission of zoonotic viruses. In contrast, the reemergence of H1N1 influenza A virus in 1977 showed a change in selection, consistent with the hypothesis of passage in a laboratory setting prior to its reintroduction into the human population, purportedly during a vaccine trial. Holistic phylogenetic analysis of selection regimes can be used to detect evolutionary signals of host switching or laboratory passage, providing insight into the circumstances of past and future viral emergence.

## INTRODUCTION

Human history has been marked by viral epidemics of zoonotic origin—viruses transmitted from a vertebrate reservoir to humans—recently including Ebola virus, Marburg virus, HIV-1, influenza A virus, SARS-CoV, MERS-CoV, SARS-CoV-2, and monkeypox virus ^1–8^. The processes by which a zoonotic spillover initiates a human epidemic are unobserved and unclear. A traditional framework of human pathogen emergence posits that circulating zoonotic viruses do not possess the ability to sustain human-to-human transmission and must evolve necessary traits to spread and persist in humans ^9,10^. This adaptation is predicted to occur before sustained human outbreaks, either in the natural host reservoir, in an intermediate host, or during initial stuttering transmission in the human population. An alternative framework states that some viruses in the natural host reservoir already have the ability to sustain human-to-human transmission, and these viruses are the source of zoonotic epidemics when introduced into the human population, and subsequent viral adaptation improves transmissibility or facilitates spread ^11–13^.

Several instances of viral adaptation to increase transmissibility shortly after a novel zoonotic virus establishes in humans have been documented. During the 2002–2004 SARS outbreak in humans, the virus evolved increased binding efficiency to human ACE2, which is an essential receptor for successful infection of human cells ^14–16^. During the 2014-2016 Ebola epidemic in West Africa, the virus accumulated mutations that increased tropism for human cells while reducing tropism for the animal host cells ^17^. SARS-CoV-2 accumulated advantageous mutations within the first year of widespread circulation, including D614G which increased transmissibility ^18,19^ and N501Y, which together with a cluster of other mutations that arose in multiple variants of concern ^20^, increased receptor binding efficiency.

As viruses switch between host species or experience other changes in their environments, the intensity, genomic hotspots, and temporal dynamics of selective forces on their genome change as well. Comparative phylogenetic methods which estimate the relative rates of non-synonymous (dN) and synonymous (dS) substitution have found broad use in understanding how viruses evolve adaptations ^21–25^. The ratio of these rates, dN/dS or ω, is a useful and informative statistic describing the nature of selective forces, whereby ω < 1 indicates purifying selection, ω > 1 indicates positive diversifying selection, and ω ∼ 1 indicates neutral evolution. Modern methods for estimating ω account for spatial and temporal heterogeneity of selective pressures, and correct for many confounding processes such as recombination, variation in synonymous substitution rates, nucleotide substitution biases, and unequal codon frequencies ^26^.

Using phylogenetic methods to characterize ω, we can describe how selection changes across different environments. For example, relaxation of natural selection can be observed when viruses are passaged in cell culture (in the absence of artificial selection for particular traits), such as with live-attenuated vaccine viral strains ^27,28^. We attribute this relaxation to the virus being released from host-specific selective pressures that had operated in its previous environment. For instance, viral variants with antigen-altering mutations would no longer have an advantage in a viral population being fed a steady supply of cells in petri dishes free of antibodies and T-cells. Hence, the branch of the evolutionary tree that corresponds to the period when this change in selective regime occurred may show a decrease in positive selection (shifting a relatively high ω down closer to 1) relative to evolution in the background context. Conversely, purifying selection that had previously acted to conserve optimized traits, such as host-specific receptor binding or host-to-host transmission ability, may be absent or less potent (shifting a relatively low ω up closer to 1),signifying relaxed selection.

In contrast, when virus is passaged with artificial selection, such as passaging in cold temperatures to induce temperature sensitivity in influenza A virus ^29^, we expect an intensification of selection. This change in selection may be driven by factors such as intensifying positive selection helping viruses adapt to a new environment and intensifying purifying selection due to a limited portion of virus variants surviving the new environment.

Here, we use the RELAX ^30^ modeling framework to conduct a formal statistical comparison between sets of branches in viral phylogenies. This approach is specifically suited to determine whether viral genomes experience intensified or relaxed selection (both purifying and diversifying selection combined) when they transition to different evolutionary environments, which are represented by sets of branches in a phylogenetic tree. As recombination and reassortment in the evolutionary history of zoonotic viruses can confound selection inference, we have extended the RELAX framework to account for multiple evolutionary histories in different genomic segments.

To investigate if adaptation is needed prior to successful zoonosis, we characterized genome-wide selection regimes for zoonotic viruses (i) in their natural animal hosts, (ii) on the stem branch of the phylogenetic tree immediately preceding a zoonotic emergence, (iii) and in the early stages of outbreaks in humans. These three selection environments are represented by disjoint sets of branches in a phylogenetic tree. The stem branch that precedes the most recent common ancestor (MRCA) of an outbreak represents transmission in the early outbreak before the MRCA of sampled lineages as well as any evolution associated with unsampled intermediate hosts ^31^. If adaptation facilitating human-to-human transmission occurs prior to the zoonotic jump or during stuttering transmission, we would expect a detectable change in the selection regime on the stem branch preceding the human outbreak. Alternatively, if the ability to transmit between humans is not the result of selection in the natural host, intermediate host, or during stuttering transmission, but is rather a property of the virus in its natural host reservoir, then we would expect that the selection regime would not change substantively prior to emergence; only once the virus is transmitting during a successful outbreak in humans is change in selection expected. We estimate and compare the three different selection regimes around the time of recent human zoonosis in viral families: *Orthomyxoviridae*, *Filoviridae*, and *Coronaviridae*. We find that substantial adaptation prior to zoonosis is not detectable and thus is not necessary for the establishment of novel zoonotic outbreaks.

## RESULTS

We characterized selection prior to successful zoonotic outbreaks to investigate if adaptation is required prior to zoonosis for establishing a successful zoonotic outbreak. We considered selection regimes (referred to as “selection” henceforth) for three phases of each outbreak, using representative phylogenetic trees and partitioning branches into three sets. One set of branches represents evolution in the animal host reservoir. The second set of branches encompasses evolution during a human outbreak. The third set is a single stem branch which separates human and animal hosts and represents evolution prior to or near the time of zoonosis (**Fig 1**). The key question is whether this branch represents any adaptation prior to establishing a successful outbreak that might be present, be it in animal or human hosts, allowing us to explore if adaptation is needed to enable sustained human-human transmission.

**Fig 1.**
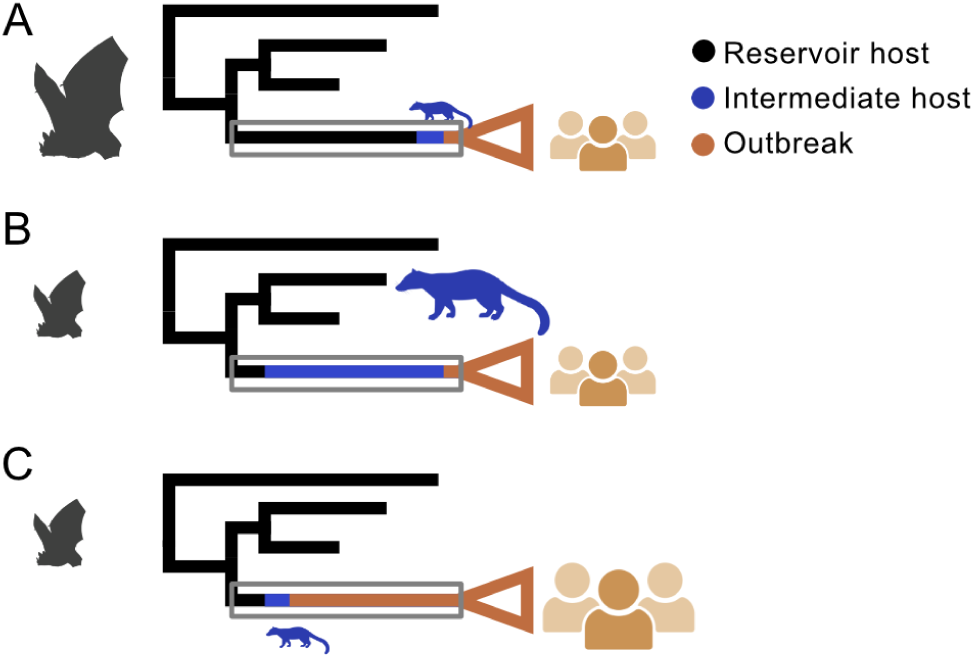
Cartoon viral phylogeny during evolutionary history of host switching. (A) Evolution on the stem leading the MRCA of human outbreak is primarily associated with the natural host reservoir. (B) Evolution on the stem leading the MRCA of human outbreak is primarily associated with (unsampled) intermediate host. (C) Evolution on the stem leading the MRCA of human outbreak is primarily associated with initial stuttering transmission in the human population, not captured by observed samples. Branches are colored by the host of the virus, the natural host reservoir (black), intermediate host (blue) or human (brown). The size of the silhouettes (bat, intermediate host, humans) reflects the proportion of the evolutionary change on the stem branch that occurred in that host. Icons created with BioRender.

We estimated and compared selection regimes using the RELAX framework ^30^. This framework captures temporal (branch to branch) and spatial (site to site in the genome) variation in selection regime using a discrete distribution of ω categories with purifying/negative (ω≤1), or diversifying/positive selection (ω≥1), using 2 or 3 categories to represent the full selection regime. The reference distribution is estimated using maximum likelihood (ML) from the “background” branches, representing evolution in the animal host reservoir, using a random effects phylogenetic likelihood framework. Selection in another set of branches (test), which can be human outbreak only, stem only, or combined (stem + human), is defined as a transformation of the reference distribution encoded as ω^K^. K is a relaxation/intensification parameter (estimated by ML). When K < 1, all the ω values in the test branches are closer to 1 (neutrality), implying that selection is overall relaxed, relative to the reference (animal host) branches. When K > 1, ω values are further away from one, implying that selection is further away from neutrality, or is intensified relative to the background. The scalar parameter K offers a convenient genome-wide measure of how selection prior to or following a zoonosis compares in intensity to selection occurring in natural reservoir hosts. Branches that are neither test nor background are treated as a nuisance set, with their own inferred ω distribution, which is not of interest. A hypothesis test where the null is K=1 (selection is identical between the environments) provides a measure of statistical significance for a change in selection.

We first examined selection dynamics preceding the zoonotic emergences of the 2009 H1N1 influenza pandemic, the 2014-2016 West African Ebola virus disease (Ebola) epidemic, and the 2004-2005 Angolan Marburg epidemic (**Fig 2A-C**).

**Fig 2.**
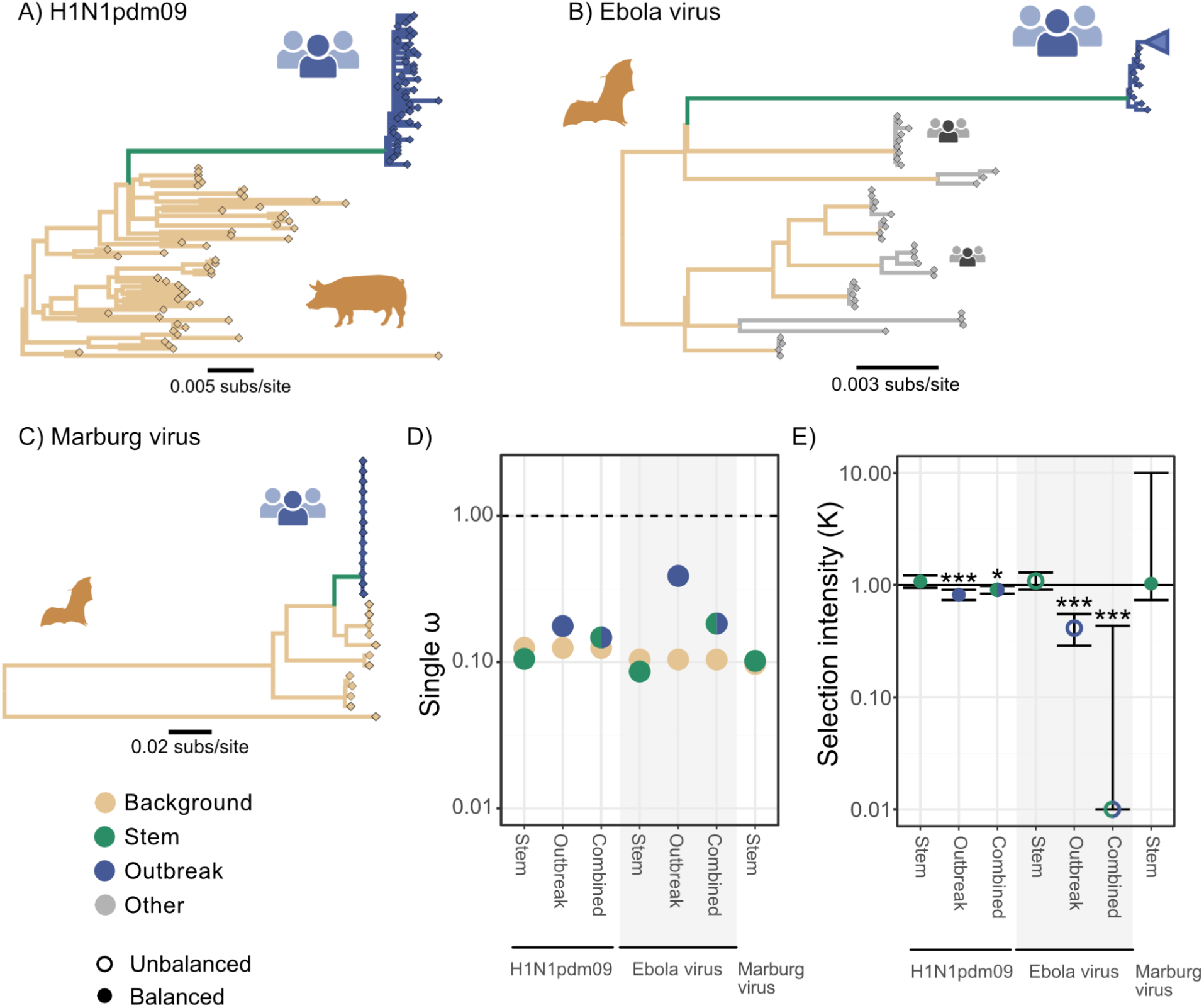
Quantifying selection regimes of key epidemics with zoonotic origin. Phylogenetic tree of (A) H1N1 2009 influenza pandemic (H1N1pdm09) (B) 2014-2016 West African Ebola outbreak (C) the Angolan 2004-2005 outbreak of Marburg virus. (D) Single ω for each branch partition of background and the test sets including “Combined” which collates the stem and the epidemic clade in test branches. Points are color coded according to the branch set: Background (brown), Stem (green), Outbreak (blue). (E) Change in selection intensity comparing selection regimes between background and test branches. Filled circles indicate a balanced model where directionality is identifiable, open circles indicate an unbalanced model and the direction above or below 1 is not identifiable (see text). For Marburg virus, there was insufficient sequence diversity within human isolates to perform the corresponding tests. Significance indicated with * *p*<0.05; ** *p*<0.01; and *** *p*<0.001. Numerical values in Supp Table 1. Icons created with BioRender.

### Selection regimes preceding and during the 2009 H1N1 influenza pandemic

The 2009 H1N1 influenza (H1N1pdm09) pandemic arose from a novel reassortment of influenza genomic segments from swine and avian viruses ^32^. The stem preceding this pandemic encompasses the estimated 9-12 years of evolution separating the epidemic most recent common ancestor (MRCA) of H1N1pdm09 in humans and their closest swine virus relatives ^32^. We found no evidence for intensification or relaxation of selection (K=1.1; *p*=0.21) on the stem branch leading to the human pandemic viruses (stem branch; **Fig 2A**) compared with the branches associated with swine viruses (reference branches; **Fig 2A**). In other words, selection on the branch preceding the emergence of H1N1pdm09 is not statistically distinguishable from typical swine influenza virus.

In contrast, we inferred relaxation of selection for the human pandemic within a year following emergence (outbreak branches; **Fig 2A**), compared with selection in swine hosts (K=0.82; *p*<0.01). During these early months of the 2009 H1N1 influenza pandemic, the single inferred ω of the virus in humans was greater than the single ω in the swine host (**Fig 2D**). This increase in ω was not driven by an intensification of positive selection but a relaxation of purifying selection. This relaxation of selection is evidence that H1N1 during the human outbreak is evolving under a different selection regime than when in swine hosts. When we include the stem together with the human branches, the relaxation of selection compared with background swine virus selection is still detectable, though less pronounced (K=0.91, *p*=0.01).

### Selection regime preceding and during filovirus epidemics

The 2014-2016 West African Ebola epidemic was the largest recorded outbreak of Ebola virus disease ^1^, occurring thousands of kilometers from previously documented human cases of Ebola virus infection. We tested for a change in the selection regime along the stem leading to the West African Ebola outbreak relative to viruses in the (presumed) bat reservoir along phylogenetic branches between human outbreaks ^33,34^ (**Fig 2B**). We exclude evolution associated with the other human outbreaks by placing outbreak clades in the nuisance set. We found no support for intensification or relaxation of selection preceding the West African epidemic (K=1.09, *p*=0.34), indicating that selection on the stem branch is consistent with selection in the bat reservoir up until the MRCA of the human epidemic clade.

There was, however, evidence for a significant change in the selection regime during the first 18 months of human-to-human transmission during the West African Ebola epidemic compared with selection on viruses in the host reservoir (**Fig 2E**; K<0.01, *p*<0.01). This signal was still detectable when we combined the stem and branches within the human outbreak (**Fig 2E**; K=0.41, *p*<0.01), though the signal was weaker. The ω distribution inferred for the reservoir/host branches only included categories of ω≤1, representing purifying and neutral selection. This ω distribution did not include a non-trivial (weight > 0) component for positive selection (ω>1) indicating that, for these Ebola virus branches, selection in the animal hosts is adequately described by purifying and neutral selection alone. This type of distribution makes it impossible to attribute the inferred change of selection in the test to a specific direction of change in the selection regime (i.e., intensification or relaxation of selection). In the situation of no ω>1 in the reference partition, an increase in positive selection (ω>1 intensifies beyond ω=1) or relaxation of purifying selection (ω≤1 relaxes to ω=1) in the test partition can only be described as a relaxation of the ω≤1. We refer to this behavior, where we observe a significant change in selection regime but cannot distinguish intensification from relaxation, as an “unbalanced model”.

Even though the RELAX analysis of Ebola virus produces an unbalanced model, the overall picture is similar to H1N1pdm09. When we compare the ω estimates from Ebola virus in humans to the bat reservoir, which are inferred with simple low resolution models that do not permit the strength of selection to vary across the genome or within clades, the epidemic clade had a higher estimate of ω, as was the case for H1N1pdm09.

Like Ebola virus, Marburg virus is a filovirus associated with hemorrhagic fever. Unlike Ebola virus bats are known to serve as the natural host reservoir of Marburg virus ^35^. One of the largest known Marburg outbreaks in humans occurred in Angola in 2004-2005 ^8^. We characterized selection along the stem preceding the epidemic MRCA and compared it with selection on branches associated with Marburg virus in bats (**Fig 2C**).There was no evidence of intensification or relaxation on the stem (K=1.03, *p*=0.90). We were unable to estimate selection regimes within the human Marburg outbreak due to insufficient viral genomic diversity among the human samples.

### Selection regimes in Betacoronaviruses prior to spillovers in humans

Betacoronaviruses are a genera of viruses circulating in bats that have repeatedly jumped into humans. SARS-CoV and SARS-CoV-2 are Betacoronavirus of the subgenus Sarbecovirus. The 2002-2004 SARS epidemic was traced back to palm civets (*Paguma larvata*) and other animals sold for human consumption in live-animal markets in China ^36^. The close contact between humans and these CoV-infected intermediate host animals likely facilitated repeated spillovers. To understand the emergence of SARS-CoV, we analyzed 139 SARS-CoV-like sarbecovirus genomes sampled in bats, two human SARS-CoV genomes representing distinct zoonotic introductions, and one SARS-CoV genome sampled from a palm civet intermediate host (**Fig 3A**). Using coding sequences across 31 putatively non-recombinant genomic regions, we compared selection along the stem leading from bat SARS-CoV-like viruses to the MRCA of human and palm civet SARS-CoV. We found no significant difference in selection between the stem and animal hosts (**Fig 3C**; K=0.97, *p*=0.91). The lack of change suggests that, prior to the MRCA of non-bat virus, the SARS-CoV-like virus had not experienced extensive evolution in an evolutionary environment different from the bat host. However, when we include the clade comprising palm civet and early human cases—primarily representing evolution in the intermediate palm civet host preceding introduction into humans—we find that the SARS-CoV selection regime was significantly relaxed compared with related bat viruses (K=0.65, *p*<0.01). The overall single inferred ω increased due to relaxation of purifying selection (rather than an increase in ω primarily due to an increase in positive selection). This relaxation in selection is evidence that after the host switch the virus is in a new evolutionary environment.

**Fig 3.**
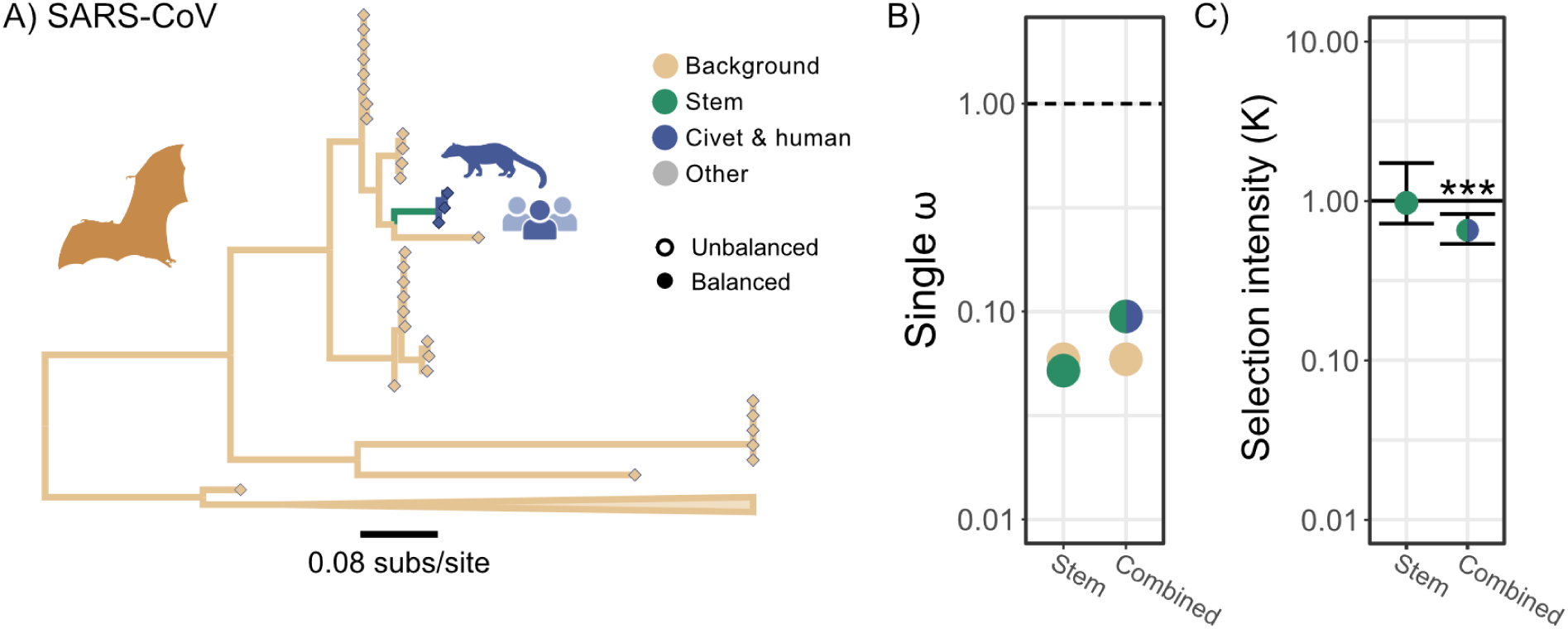
Quantifying selection regime in SARS-CoV. **(A)** Phylogenetic tree of SARS-CoV like sarbecoviruses non-recombinant region 22, branch color indicate partition. The SARS outbreak includes 2 human sequences and 1 civet sequence, which are monophyletic across all regions. (B) Single ω for each branch partition of branch sets. Points are color coded according to the branch set: Background (brown), Stem (green), or Outbreak (blue). (C) Change in selection intensity between background and test sets. Filled circles indicate a balanced model where directionality is identifiable, open circles indicate an unbalanced model and the direction above or below 1 is not identifiable. Numerical values in Supp Table 1. Significance indicated with * *p*<0.05; ** *p*<0.01; and *** *p*<0.001. Icons created with BioRender.

The circumstances surrounding the emergence of SARS-CoV-2 are similar to SARS-CoV. In late 2019, SARS-CoV-2 began spreading at a market selling wide animals in central China ^4,37^, thousands of kilometers away from closely related bat viruses in southern China and Laos ^38^. Genetic and epidemiological evidence supports the hypothesis that virus preceding SARS-CoV-2 briefly circulated in an intermediate host sold at the market ^39–42^. However, it has been suggested that SARS-CoV-2 was passaged or modified in a laboratory context, prior to emergence ^43^.

If there was extensive evolution in an intermediate host or passage in a laboratory context prior to emergence, we would expect detectable change in selection on the stem preceding SARS-CoV-2. However, our analysis of selection on the stem preceding SARS-CoV-2 emergence across 15 putatively non-recombinant regions found no evidence of intensification or relaxation of selection compared with selection of the bat host reservoir (K=1.1, *p*=0.23; **Fig 4**). Hence, we find no evidence to suggest SARS-CoV-2 experienced selective pressure in an environment different from related bat viruses prior to its emergence in humans. This result does not change if we use a different approach to identifying non-recombinant regions (K=1.02, *p*=0.82; **Supp Fig 1**), and is consistent with previous analysis ^22^.

**Fig 4.**
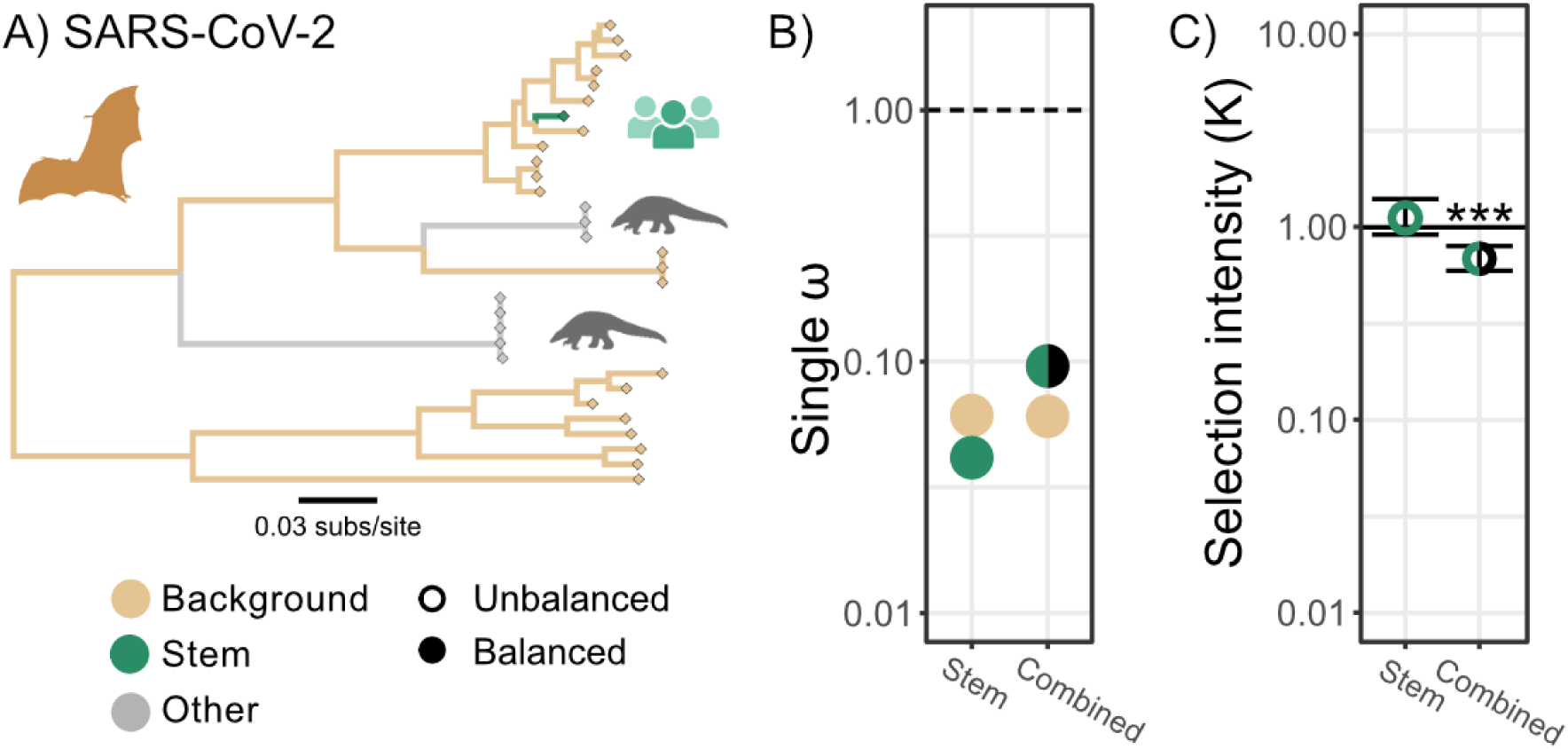
Quantifying selection regimes in SARS-CoV-2. (A) Phylogenetic tree of SARS-CoV-2 like sarbecoviruses non recombinant region 08, branch color indicate partition. (B) Single ω for each branch partition of background and the test sets and includes both the stem and outbreak partitions in the test (**Supp Fig 2**). Points are color coded according to the branch set: Background (brown), Stem (green), or Outbreak (black). (C) Change in selection intensity comparing selection associated with background to test set. Filled circles indicate a balanced model where directionality is identifiable, open circles indicate an unbalanced model and the direction above or below 1 is not identifiable. Numerical values in Supp Table 1. Significance indicated with * *p*<0.05; ** *p*<0.01; and *** *p*<0.001. Icons created with BioRender.

We then examined evolution along the SARS-CoV-2 stem in combination with viral evolution during the first 3 months of the outbreak in China, to understand the selection environment of SARS-CoV-2 in humans compared with the bat host reservoir. We find evidence for a significant change in selection regime, consistent with a host switch causing a change in the evolutionary environment (K=0.69, *p*<0.01), as previously described ^22^. As with the West African Ebola epidemic we cannot confidently infer the directionality of this change because the model is unbalanced. The change in selection regime is also detectable when the clade of the first world-wide wave, up to October 2020, is included with the stem compared with bat virus background (K=0.56, *p*<0.01; **Supp Fig 1**).

### The reemergence of H1N1 in 1977

In 1977, H1N1 influenza A virus reemerged in humans after going extinct 20 years prior. The reemergent virus was closely related to a strain that had been circulating in the 1950s ^44,45^. It has been proposed that H1N1 reemerged in 1977 without expected evolution because it was frozen in a laboratory before itwas accidentally allowed to re-establish “wild” human-to-human transmission, perhaps involving (i) a live-attenuated vaccine virus or (ii) a laboratory virus used to infect humans in a human challenge trial of an influenza vaccine ^46^. We infer only 48 substitutions across the entire genome along the stem preceding the 1977 reemergence, with multiple genomic segments experiencing only 2 substitutions (Supp Table 2). Hence, the 1977 strain is likely too genetically similar to 1950s H1N1 virus to support a natural reemergence. However, the provenance of this virus prior to reemergence remains poorly understood.

We compared the selection regime of the coding regions across all H1N1 influenza A virus segments on the stem preceding the 1977 influenza pandemic with evolution post-1977 (**Fig 5**). We found evidence for significant relaxation of selection along the stem (K=0.71, *p*=0.043), in addition to an increase in ω (**Fig 5C**). This change in selection regime suggests that in addition to being frozen between the 1950s and 1977, the precursor to the 1977 H1N1 influenza pandemic experienced a limited amount of evolution in an environment distinct from human-to-human transmission (e.g. laboratory passage for attenuation or under artificial selection). This result remains supported even if we exclude the phylogenetic tips associated with influenza A virus isolates that have a known history of laboratory passaging (**Supp Fig 2**; K=0.70, *p*=0.042).

**Fig 5.**
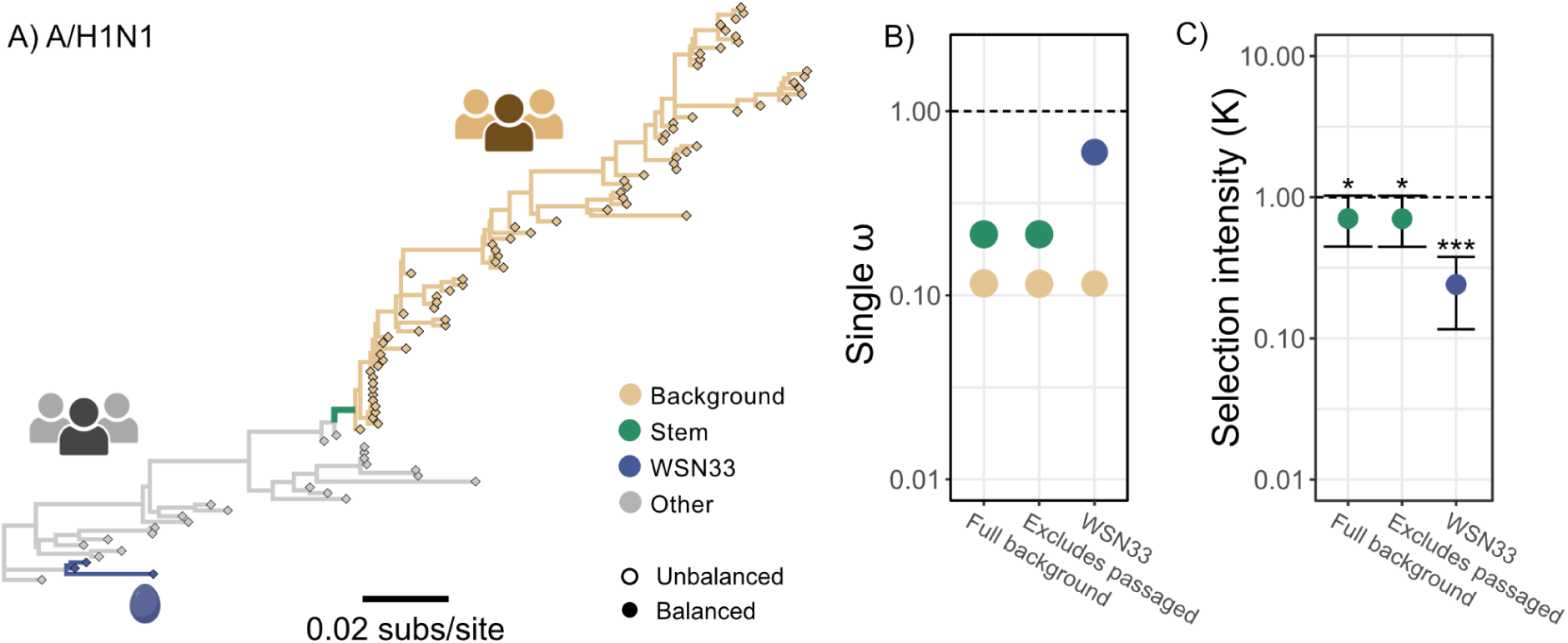
Quantifying selection regime of H1N1. **(A)** Phylogenetic tree of H1N1 HA segment, branch color indicates partition. The stem branch connects 1977 clade and basal ancestors, and is foreground for 1977 flu test. WSN33 (blue) branches are clade associated with laboratory passage. Other branches are excluded from analysis (treated as a nuisance group). (B) Single ω for each branch partition of background and the test sets. “Full background” compares selection on stem to displayed background. “Excluded passaged” compares selection on Stem to background excluding tips which have history of passaging (49/74; **Supp Fig 3**) “WSN33” compares selection of WSN33 branches to the full background. Points are color coded according to the branch set: Background (brown), Stem of 1977 pandemic (green), or WSN33 clade (blue). (C) Change in selection intensity comparing selection associated with background to test set. Filled circles indicate a balanced model where directionality is identifiable, open circles indicate an unbalanced model and the direction above or below 1 is not identifiable. Significance indicated with * *p*<0.05; ** *p*<0.01; and *** *p*<0.001. Numerical values in Supp Table 1. Icons created with BioRender.

### Laboratory passage of H1N1 influenza A viruses

To better contextualize the atypical evolutionary dynamics preceding the 1977 H1N1 influenza pandemic, we explored whether the signature of laboratory passage would be detected as a change in the selection regime by RELAX. The Wilson Smith 1933 H1N1 (WSN33) isolate is the ancestor of many laboratory strains used in research. We compared the selection regime of 3 early laboratory isolates derived from WSN33 with the selection regime associated with human-to-human H1N1 transmission across all 8 genomic segments (**Fig 5**). There was significant evidence of relaxation of selection in the WSN33 clade (K=0.24, *p*<0.01; **Fig 6**). This relaxation of selection may be due to the removal of host-specific constraints, such as immune evasion driving positive selection, and maintenance of host-optimized receptor binding driving purifying selection.

**Fig 6.**
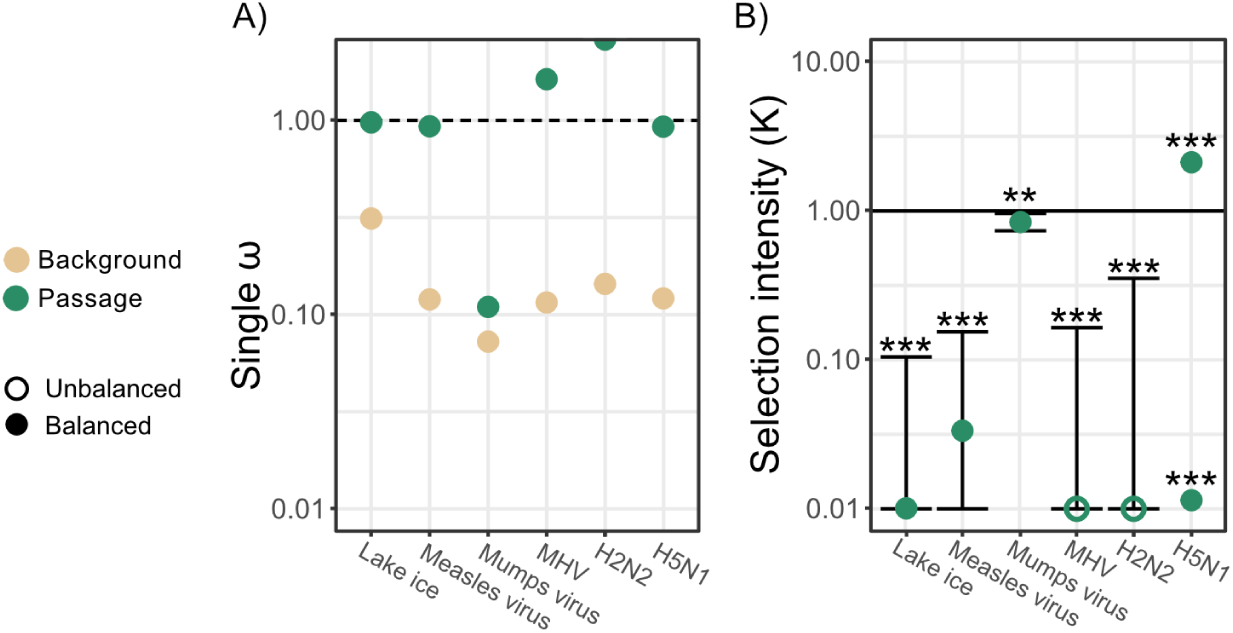
Quantifying selection regimes of laboratory passaged viruses. **(A)** Single ω for each branch partition of background and the test sets. Points are color coded according to the branch set: Background (brown), and Passage (green). (B) Change in selection intensity, comparing selection associated with transmission between humans (except H5N1 which is compared with selection in avian host reservoir, and MHV to selection associated with murine host) to i) positive control of H1N1 (Lake ice) ii) passaging in cell lines of Measles virus iii) Mumps virus and iv) Mouse hepatitis virus (MHV) v) artificial selection for cold adaptation (H2N2) and vi) gain of function passaging in ferrets to introduce airborne transmission (H5N1), points above and below K=1 indicate the instability in model estimate. Filled circles indicate a balanced model where directionality is identifiable, open circles indicate an unbalanced model and the direction above or below 1 is not identifiable. Significance indicated with * *p*<0.05; ** *p*<0.01; and *** *p*<0.001. Numerical values in Supp Table 1. Trees for analysis in Supp Fig 3-8. Icons created with BioRender.

Zhang et al., ^47^ claimed that influenza A virus persisted in lake ice based on HA gene sequences. However, Worobey ^48^ noted (i) that these putative lake ice viruses were monophyletic and descended from the WSN33-derived positive control and (ii) that they had evolved greatly from their common ancestor (**Supp Fig 3**). This finding indicated that these samples were contaminants derived from this ab strain ^48^, which had in fact been used as a positive control by Zhang et al ^47^. Accordingly, in the current study, we found that selection on putative lake ice viruses was significantly relaxed compared with human H1N1 virus (K=0.1, *p*<0.01; **Fig 5**), which is consistent with laboratory passaging in the absence of typical selective forces. Clearly, these were “lab”, not “lake”, viruses.

### Selection on attenuated vaccine viruses

Measles virus and mumps virus passaged in cell culture to create attenuated viruses for vaccines were found to have evolved with increased ω, consistent with a relaxation of purifying selection ^27,28^. We compared the selection regime of measles virus and mumps virus viruses passaged for vaccine attenuation with selection on branches representing transmissions between humans (**Supp Fig 4; Supp Fig 5**). In agreement with previous results, the vaccine attenuated measles virus and mumps virus both have an increased ω relative to the background branches (**Fig 6A**). Furthermore, we were able to detect significant relaxation of selection in the vaccine attenuated viruses relative to selection in the human host for both measles virus (K=0.03, *p*<0.01) and mumps virus (K=0.84, *p*<0.01) (**Fig 6**).

### Serially-passaged coronavirus

Murine hepatitis virus (MHV) is a betacoronavirus naturally found in mice. Numerous isolates of MHV have been propagated in laboratory cell culture. We compared the selection regime of MHV propagated in cell culture with the selection regime of naturally transmitting MHV (**Supp Fig 6**), across 11 putatively non-recombinant regions. We inferred an increased ω in the passaged virus relative to the background. Further, we found evidence of a change in selection in the passaged viruses (K=0, *p*<0.01; **Fig 6**); however, as with the West African Ebola outbreak we are unable to identify the direction of change due to an unbalanced model.

### Artificial selection on influenza A viruses

Artificial selection, e.g., passaging of viruses under selective constraints to produce novel phenotypes, is expected to alter the selection regime. An H2N2 influenza A virus isolate was passaged at colder temperatures to attenuate virus replication at human body temperature ^29^. We compared the selection regime associated with this cold adapted virus, with selection associated with H2N2 virus transmitted between humans (**Supp Fig 7**). We found evidence of a change in selection of the non-surface proteins resulting from artificial selection (K=0, *p*<0.01; **Fig 6**), however the model is unbalanced and we cannot identify the direction of selection intensity change. Notably, we also inferred an increase in ω in the passaged viruses relative to the virus in humans, consistent with adaptation due to laboratory passage.

In an artificial selection experiment, H5N1 influenza A virus was passaged between ferrets to gain airborne transmissibility ^49^. We inferred a significant change in the selection regime due to this artificial selection in the new host compared with virus in the host avian reservoir (**Fig 6; Supp Fig 8**; *p*<0.01). However the inferred direction of change in selection intensity was unstable; relaxation (K=0.01) and intensification (K=2.12) did not have significantly different likelihoods in our model; this instability is partly due to the relatively low diversity of available sequence. Across our analyses of viruses passaged in a laboratory setting, we consistently found evidence for a change in the selection regime of passaged viruses, with and without artificial selection, compared with evolution in the host environment.

## Discussion

How much adaptation must a zoonotic virus undergo before it becomes adept at sustained human-to-human transmission and where does this adaptation take place? Observed empirical data from early during outbreaks can be sparse and inconclusive; however, we can infer evolutionary patterns using a phylogenetic framework. We reason that sufficiently extensive adaptation prior to an outbreak in humans will leave detectable evolutionary signals in the viral genome, specifically on the phylogenetic branch immediately preceding the observed outbreak. We also identified several instances of viral passaging in culture or animals that provide “controls” for our statistical framework.

We find that for four recent zoonotic epidemics—H1N1pdm2009, Ebola virus, Marburg virus, and SARS-CoV-2—these viruses continued to evolve in a manner indistinguishable from that in their host reservoir up to spillover into humans. These findings challenge the model where zoonotic viruses must progressively evolve the ability to sustain human-to-human transmission ^9,10,50^. Instead, this evolutionary pattern is consistent with a process whereby humans are repeatedly exposed to viruses originating from host reservoirs, but only viruses that happen to be sufficiently transmissible in humans to sustain transmission tend to be the seed of zoonotic epidemics in the human population ^11,12^. In contrast, we find evidence of a change in selection regime preceding the emergence of SARS-CoV in humans and the reemergence of influenza A/H1N1 in 1977, suggesting that in these cases virus evolved under a different evolutionary environment, respectively in an intermediate host and laboratory passaging, prior to emergence.

Previous analysis of SARS-CoV-2 and its closest relatives found little evidence of significant genetic adaptation favoring human-to-human transmission during the evolutionary period immediately preceding the human outbreak ^22^. Our results are consistent with this conclusion. We did not detect any significant change in the selection regime on the pre-outbreak stem compared with the selection regime in bats, change which would be expected had the virus been extensively passaged in a laboratory setting or had it been circulating in an intermediate host for a prolonged period. If, as a large body of evidence indicates, SARS-CoV-2 did circulate in an intermediate mammalian host before jumping into humans ^39–42^, the duration of this circulation must have been brief ^41^ and was not sufficient to leave a sign of a change in the selective regime.

We found that selection on the branches leading to early human SARS-CoV genomes was detectably different from selection in the natural bat host reservoir. We attribute this change in selection to diversity accumulated in an intermediate host: palm civets ^36^, consistent with adaptation within an intermediate host (**Fig 1B**). Although we detected a change in selection regime in palm civets, mutations associated with improved transmission in humans were acquired later during transmission between humans, and not during circulation within palm civets ^14–16^. This observation stands in contrast to SARS-CoV-2, where the change in selection regime associated with an intermediate host was not detectable. The lack of detectable selection prior to the MRCA of SARS-CoV in palm civets and humans is similar to the evolutionary dynamics prior to the emergence of SARS-CoV-2 in humans, consistent with a lack of a change in the selection regime in bats prior to host switching.

Bats are thought to serve as the host reservoir of not just sarbecoviruses, but also Ebola virus. The appearance of a Central African strain of Ebola virus in humans in West Africa in 2014 was unexpected ^51–53^, raising questions about how the virus traveled so far without being previously detected. However, there is serological evidence that people in West Africa have been exposed to Ebola virus prior to 2013 ^54^, suggesting that there had been repeated introductions of Ebola virus to humans in the region without sustained transmission for years prior to the epidemic. However, these dead-end introductions cannot exert selective pressure on the virus in the bat reservoir. Correspondingly, we find no evidence of change in selection prior to Ebola virus emergence of the 2014-2016 outbreak, suggesting Ebola virus remained in the natural host until it emerged in the human population and sustained transmission.

Unlike the previous examples of zoonotic epidemics, there is evidence of a change in selection preceding the reemergence of influenza A/H1N1 virus in 1977, which provides a new line of evidence supporting the theory that its reemergence was associated with an influenza vaccine challenge trial ^46^. If the precursor to the 1977 H1N1 influenza pandemic had been frozen for over two decades prior to its reemergence, and not passaged, the selection regime along the phylogenetic branches leading to the re-emergent virus would be similar to that of other circulating H1N1 viruses. Instead, we found that the virus preceding the 1977 influenza pandemic evolved under a selection regime markedly different from typical human-to-human transmission; this change in selection is consistent with the change in selection of influenza A virus during laboratory passage (**Fig 5-6**). Further, it has been previously noted that viral isolates sampled shortly after the 1977 reemergence were temperature sensitive, and cold adaptation is a trait that is frequently artificially selected for in laboratory passage to make live attenuated influenza vaccines ^29,55^. These observations of evolutionary stasis, temperature sensitivity, and change in selection, strongly suggest that the precursor to 1977 H1N1 was both frozen and passaged prior to reemergence, and its introduction into the human population is consistent with an escape during live attenuated influenza vaccine trial. Accordingly, the 1977 H1N1 virus and its descendants could be designated a putative circulating vaccine-derived influenza A virus (“cVDIAV”), following the convention used when a similar process occurs with live-attenuated oral poliovirus vaccines (i.e., circulating vaccine-derived polioviruses, “cVDPVs”).

Our approach to characterizing selection regimes overcomes a common shortcoming of similar investigations: the need to independently analyze non-recombinant or non-reassortant genomic segments ^30,56,57^. We extended the RELAX phylogenetic framework to account for different evolutionary histories by applying a single selection regime model to multiple genomic segments with distinct phylogenetic histories. We applied this multi-region analysis to combine regions for viruses where recombination or reassortment were likely (e.g., positive sense RNA viruses and influenza A virus, respectively). Using multiple genomic regions in the same model increased the power to distinguish changes in the selection regime.

In this study we have distinguished epidemics that are characterized by viruses that evolved primarily under a selection regime in the natural host reservoir prior to emergence from those that did not. Humans are constantly exposed to animal viruses ^54,58^. However, most of these exposures do not result in ongoing outbreaks with human-to-human transmission, due to low fitness of the virus or lack of sufficient transmission opportunities (such as in rural communities)^31^. In recent zoonotic epidemics where there was sustained human-to-human transmission, we found no detectable change in selection preceding zoonotic emergence, suggesting that virus from the natural host reservoir is introduced into the human population and can successfully transmit without requiring prior adaptation. Applying multi-region RELAX to the stem of novel viral outbreaks is a tool that we will be able to apply to future outbreaks, to rapidly assess the possibility of evolution in an intermediate host or laboratory setting compared to zoonosis directly from the natural host reservoir.

## Methods

### Data and code availability

Accession numbers and sequence names for all data are available in **Supp** **Table 3**. Code for RELAX v2.5.62 available at [GitHub](https://github.com/veg/hyphy/releases/tag/2.5.62).

### Phylogenic selection analysis

For all influenza A virus data sets, coding regions were extracted from each genomic segment. Then each region was aligned using MAFFT (v7.409) ^59^; alignments were checked manually. For all other data sets, full genomes were aligned to the reference genome and split into non-recombinant regions. For MHV, a positive-sense RNA virus, putative non-recombinant regions were identified using GARD ^60^. For SARS-CoV and SARS-CoV-2, predefined putative non-recombinant regions were used ^61^. For all data sets, coding regions were extracted based on the reference genome annotation. For each region, a maximum likelihood phylogeny was inferred with IQtree2 (v2.0.4) ^62^ using the GTR+F+Γ_4_ model.

Change in intensity of selection was tested across all genomic regions with the multi-region (joint) RELAX model implemented in HyPhy (v2.5.62) ^30^. In the RELAX testing framework, we partition branches in a phylogenetic tree into “*test*” and “*reference*”, with an optional “*nuisance*” set for the remaining (if any) branches. Evolution along each tree branch is described using a continuous time discrete space Markov model of codon evolution (for a review see ^63^); the ratio of non-synonymous to synonymous substitution rates (ω) is used for model selection. Along each branch, the evolutionary process is modeled as an independent draw from discrete distributions on ω, with one rate allocated to positive selection (ω≥1) and one or two rates allocated to negative selection or neutral evolution (ω≤1). Phylogenetic likelihood is computed by integrating over the assignments of ω to branches, and all model parameters, including ω distributions, are estimated by maximum likelihood. The selection regimes (ω distributions) of the test and reference branches are related by a selection intensity parameter K, via ω(test) = ω(background)^K^. Selection intensity (K) greater than 1 indicates intensification, where positive and purifying increase in strength (move away from ω = 1), and K less than 1 indicates relaxation, where positive and purifying selection decrease in strength (move closer to ω=1). We use the likelihood ratio test to determine whether K is significantly different from 1 (K=1 when the selection regime in test and reference are equal).

The joint RELAX test is an extension of RELAX in which multiple evolutionary histories (i.e., trees) share the inferred selection regime (ω distributions) and selection intensity test parameter (K). We ran joint RELAX for each context with every combination of (i) with and without multiple hits, which allows instantaneous multi-nucleotide substitutions, (ii) with and without synonymous rate variation, which accommodates site-to-site variation of synonymous substitution rates, and (iii) with 2 or 3 rate partitions, which has 1 or 2 partitions for purifying or neutral selection respectively. From these 8 options, the best model was chosen with AIC.

### Data sets

#### Selection regime preceding and during the H1N1 2009 pandemic (H1N1pdm09)

To characterize the selection regime preceding H1N1pdm09, we used previously curated virus genomes ^64^, with up to 58 stains for each of the 8 influenza genome segments. Additionally, to characterize the selection regime during the initial human outbreak, we considered 39 stains selected from Influenza Virus Resource search with parameters of pandemic H1N1 virus that are full length with all segments sampled from a human host in North America from the FLU project collected from 1 April 2009 through 1 March 2010. We used branches associated with transmission between swine viruses, to represent selection associated with the host reservoir (reference partition). The stem branch leading to the MRCA of H1N1pdm09 was used as the test branch of interest. Additionally, selection on the stem branch preceding H1N1pdm09 and branches within the outbreak clade, was tested against the selection regime of the same reference partition.

#### Selection regime preceding and during Ebolavirus 2014-2016 West Africa Ebola outbreak

We assembled a dataset of 101 full length Ebola virus genomes: 61 genomes were from the West African Ebola epidemic (collected 1 January 2014 through 1 June 2015) and the remaining 40 were representative of all previous known Ebola Zaire outbreaks in humans during which a viral genome had been sequenced. All coding regions concatenated in a single open-reading frame, excluding the sections of the glycoprotein gene containing overlapping reading frames.

We identified branches between human Ebola outbreaks, to represent selection associated with the natural host reservoir (reference partition). Branches that were associated within human outbreak clades were excluded. The long branch leading to the MRCA of the 2014-2016 West African outbreak was used as the test branch of interest. Additionally, selection on the stem branch preceding the 2014-2016 outbreak and branches within the outbreak clade, was tested against the selection regime of the same reference partition.

#### Selection regime preceding and during Marburg virus disease Angolan outbreak

We analyzed all Marburg virus genomes sampled from bats available on NCBI (n=12) and 14 sampled from the 2004-2005 outbreak in Angola ^8^ which is the largest known human outbreak of Marburg virus disease to date. Similarly to Ebola virus, Marburg virus is a negative-sense single-stranded RNA virus, and we concatenated all coding regions for a single genome wide analysis. The branches associated with bat samples were used to represent selection in the natural host reservoir, which is understood to be bats. The stem branch leading to the human outbreak was used as the test branch of interest.

#### Selection regimes in SARS-CoV-like virus

In 2002-2004, SARS-CoV spilled over into humans, likely from palm civets (*Paguma larvata*) sold for human consumption in live-animal markets in China ^36^. We used a total of 142 sequences, including 3 SARS-CoV sequences (2 from humans associated with recent introductions and 1 from a palm civet), using 31 predefined putative non-recombinant regions ^16,61^. We considered selection preceding the SARS outbreak by comparing the stem branch leading to the MRCA of SARS-CoV civet and human samples to the branches associated with a bat host. We also compared the early outbreak, using all branches in the SARS-CoV clade with the branch preceding it to selection in the host, using the same reference partition.

#### Selection regimes in SARS-CoV-2-like virus

We considered the selection of SARS-CoV-2-like virus preceding COVID-19 pandemic and compared it to selection of SARS-CoV-2-like sarbecoviruses sampled from the natural bat host reservoir. We analyzed a total 31 genomes ^61^ with 15 pre-defined putatively non-recombinant regions ^38^. The robustness of changes in selection to non-recombinant regions was tested by repeating the RELAX analyses with non-recombinant regions inferred by 3seq ^65^, which identified 21 regions. We considered selection preceding the COVID-19 outbreak by comparing the stem branch leading to SARS-CoV-2, represented by the reference sequence Hu-1 to branches associated with the natural host reservoir of bats.

We also considered selection early in the COVID-19 pandemic using two different contexts. First, we included a subsample of 50 SARS-CoV-2 genomes representing the early outbreak in China, over 3 months, from a previous dataset ^41^. Second, we analyzed a random sample of 39 sequences from NCBI Virus with complete genomes sampled from 15 September 2020 to 1 October 2020 to represent early worldwide transmission. For each of these sequence sets representing the early outbreak, we compared selection of the host reference partition to the outbreak clade and the stem leading to SARS-CoV-2.

Of the SARS-CoV-2-like sarbecoviruses, 8 were sampled from pangolins (the rest from bats), which form 2 pangolin clades ^66,67^. The stem leading to the pangolin clades, and the pangolin clades were excluded from the reservoir host background.

#### Selection of H1N1 influenza A viruses

In 1957 A/H1N1 was replaced by another strain of influenza A virus: H2N2. In 1977, an H1N1 strain nearly identical to that which circulated in the 1950s reemerged in the human population^44^. We used 100 genomes sampled from before and after 1977 to build a phylogenetic tree and define the branch leading to the 1977 influenza pandemic as the stem. To characterize typical human-to-human transmission, we considered the branches descended from the 1977 influenza pandemic. We excluded branches associated with samples from before 1977 from the background due to concerns of laboratory passaging, erroneous inclusion of lab strains, and sequence quality. As a robustness check, we performed an additional analysis excluding 49 tips associated with post-1977 isolates with a documented history of laboratory passaging.

We inferred the synonymous and nonsynonymous substitutions preceding the 1977 reemergence using TreeTime (v0.8.6) ^68^ ancestral state reconstruction.

We then tested if laboratory passage of virus isolates changed selection regime by comparing the background with the clade that includes 3 early laboratory isolates derived from WSN33 ^69^: WSN/33, and two derivatives A/United Kingdom/1/1933 and A/Wilson-Smith/1933(H1N1), all of which are known to be passaged before sequencing.

#### Selection of H1N1 influenza A viruses putatively from lake ice

Zhang et al.,^47^ purportedly isolated H1N1 influenza A virus from Serbian lake ice and suggested these samples were frozen in the ice from a local outbreak. We considered the 19 HA sequences purportedly isolated from lake ice, 1 positive control ^47^, and 19 HA genes from the H1N1 dataset from the 1930s through 1950s, excluding tips associated with sequences that have a history of lab passaging ^48^ for a total of 38 sequences. Branches associated with transmission between humans were used for the reference partition, and branches in the ‘lake ice’ clade were used as the test partition.

#### Selection on attenuated vaccine measles virus

We analyzed 28 measles virus full genomes that were assembled previously for a molecular clock analysis^70^. This set of genomes is characteristic of measles virus evolution within humans, and we used branches associated with these samples to characterize background selection. We then used 13 genomes from vaccine associated strains of measles virus, previously identified^28^, which were passaged in cell culture to produce an attenuated virus intended for use as a vaccine. Branches associated with the vaccine strains were used as the test branches.

#### Selection on attenuated vaccine mumps virus

We analyzed 59 mumps virus genomes that were selected by subsetting from Nextstrain (https://nextstrain.org/mumps/na), and collecting the associated full genomes from NCBI. Branches associated with these samples were used to characterize background selection. We characterized mumps virus passaged in the lab for attenuation, with 10 mumps virus genomes associated with vaccine strains, which were previously identified ^28^.

#### Serially-passaged coronavirus MHV

MHV is a betacoronavirus naturally found in mice. Graepel *et al*. passged MHV 250 times in cell culture ^71^. We used the passaging control that did not include experimental modifications to characterize selection associated with laboratory passaging, using branches leading to the passaged virus from the input wild type strain. Background selection was characterized with branches between 5 MHV lineage defining strains. 11 non-recombinant regions were inferred using GARD.

#### Artificial selection on influenza A virus A/H2N2

An H2N2 influenza A virus (wt A/Leningrad/134/57) was passaged from 34°C down to 25°C to produce the non-surface proteins for a reassortant live attenuated vaccine that that was cold adapted and so had attenuated replication at human body temperature ^29^. We characterized the selection regime of passaging with artificial selection as the branches that were descendant from the wildtype leading to isolates sequenced at passages 17 and 47, and the input wild type sequence. The branch leading to the input wild type was excluded. The background context for comparing to cold adaptation of an H2N2 stain was characterized by a random subsample of 50 full H2N2 genomes from Influenza Virus Resource, with all proteins available, excluding vaccine strains and viruses sampled from a human host.

#### Artificial selection on influenza A virus A/H5N1

Herfst *et al.*, passaged virus derived from A/Indonesia/5/2005 10 times in ferrets, which was passaged twice more in two parallel sets of transmission ^49^. We considered all branches in the clade of all available passaged sequences (n=6) and the branch leading from the initial genome (A/Indonesia/5/2005) to this clade as the test partition. The background context was characterized by a random subsample of 94 whole H5N1 genomes from Influenza Virus Resource, with all proteins available, sampled from avian sources from 1980 to 2005. We compared selection dynamics along with branches associated with the H5N1 samples from the avian population (reference set). The branch leading to the human A/Indonesia/5/2005 from the avian samples was excluded from main analysis.

## Acknowledgments

We gratefully acknowledge all the authors and contributors who generated and shared the viral genomic sequences and metadata, including the originating and submitting laboratories who shared data on GISAID (available at [EPI_SET_240827gw](https://epicov.org/epi3/epi_set/240827gw?main=true)), on which this research is based. We thank Stephen A. Goldstein, Mark Zeller, Tetyana I. Vasylyeva, and Marc Suchard for their insightful discussions.

This work was funded in part with federal funds from the National Institute of Allergy and Infectious Diseases National Institutes of Health, National Institutes of Health (NIH-NIAID) and National Science Foundation (NSF). J.L.H. acknowledges support from NIH (grant R01AI153044). S.K.L.P and J.D.Z. acknowledge support from NIH (AI183870, GM151683, GM144468) and the NSF (grant DBI/2419522). J.E.P. acknowledges support from NIH-NIAID (T15LM011271) and the UC San Diego Merkin Fellowship. M.W acknowledges support from NIH-NIAID (contract no. 75N93021C00015). E.P. and K.G.A. acknowledge support from the NIH (grant U01AI151812). K.G.A. also acknowledges support from the NIH (grant U19AI135995). J.O.W. acknowledges support from NIH-NIAID (R01AI135992).

## Author contributions

Conceptualization: J.L.H, and J.O.W.; Methodology: J.L.H., S.L.K.P., J.D.Z., and J.O.W.; Software: J.L.H., S.L.K.P., and J.D.P.; Validation: J.L.H., S.L.K.P., and J.D.Z.; Formal analysis: J.L.H., S.L.K.P., J.D.Z., and J.O.W.; Investigation: J.L.H., J.E.P., E.P., M.W., K.G.A., and J.O.W.; Resources: S.L.K.P., and J.O.W.; Data curation: J.L.H., J.E.P., E.P., M.W., K.G.A., and J.O.W.; Writing – original draft preparation: J.L.H., S.L.K.P., and J.O.W.; Writing – review and editing: All authors.; Visualization: J.L.H., S.L.K.P., J.D.Z., and J.O.W.; Supervision: S.L.K.P., K.G.A., M.W., and J.O.W.; Project administration: J.L.H., and J.O.W.; Funding acquisition: S.L.K.P., K.G.A., M.W., and J.O.W.;

---

**Supp Fig 1.**
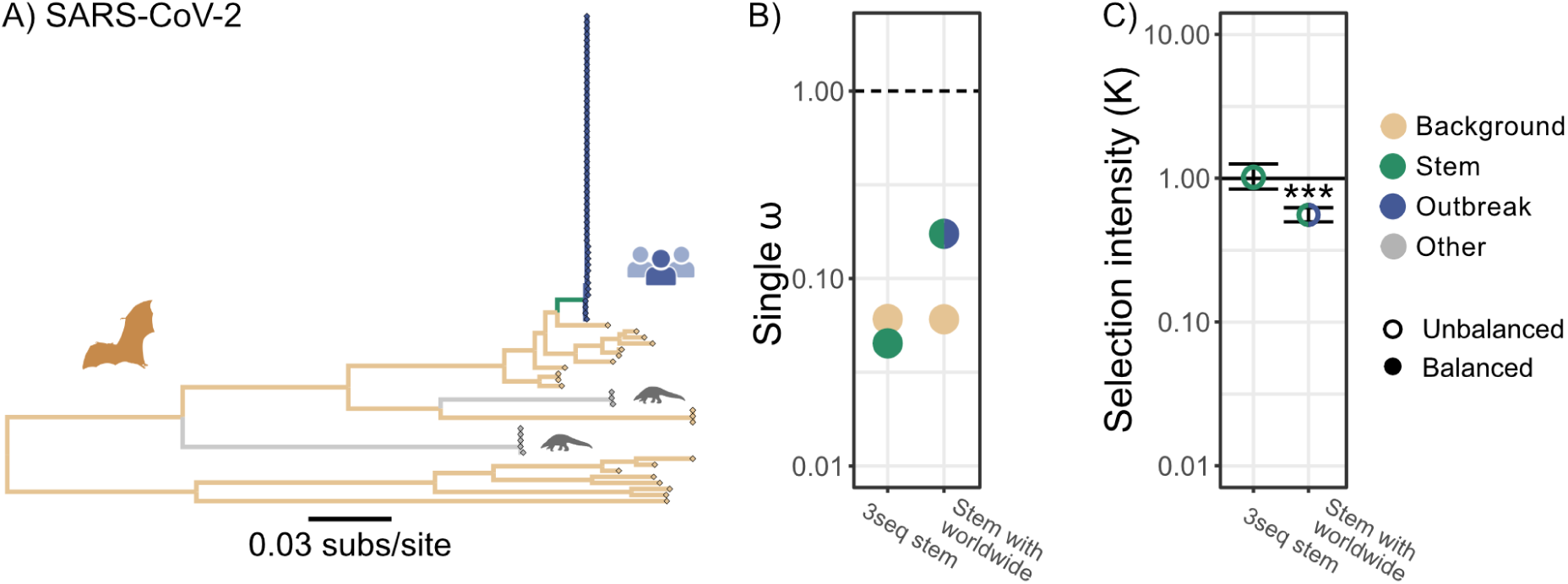
Quantifying selection regimes in SARS-CoV-2. (A) Phylogenetic tree of SARS-CoV-2 like sarbecoviruses, with SARS-CoV-2 early worldwide outbreak characterized by 39 sequences sampled up to October 2020, non recombinant region 08, branch color indicate partition. (B) Single ω for each branch partition of background and the test sets, including 3seq stem, which compares the stem branch with background, based on alignment of non-recombinant regions inferred with 3seq (no tree displayed), and “combined with worldwide” which compares stem and worldwide outbreak to background, color coded to match the tree. Points are color coded according to the branch set: Background (brown), Stem (green), or Outbreak (blue). (C) Change in selection intensity comparing selection associated with background to test set. Filled circles indicate a balanced model where directionality is identifiable, open circles indicate an unbalanced model and the direction above or below 1 is not identifiable. Numerical values in **Supp Table 1**. Significance indicated with * *p*<0.05; ** *p*<0.01; and *** *p*<0.001. Icons created with BioRender.

**Supp Fig 2.**
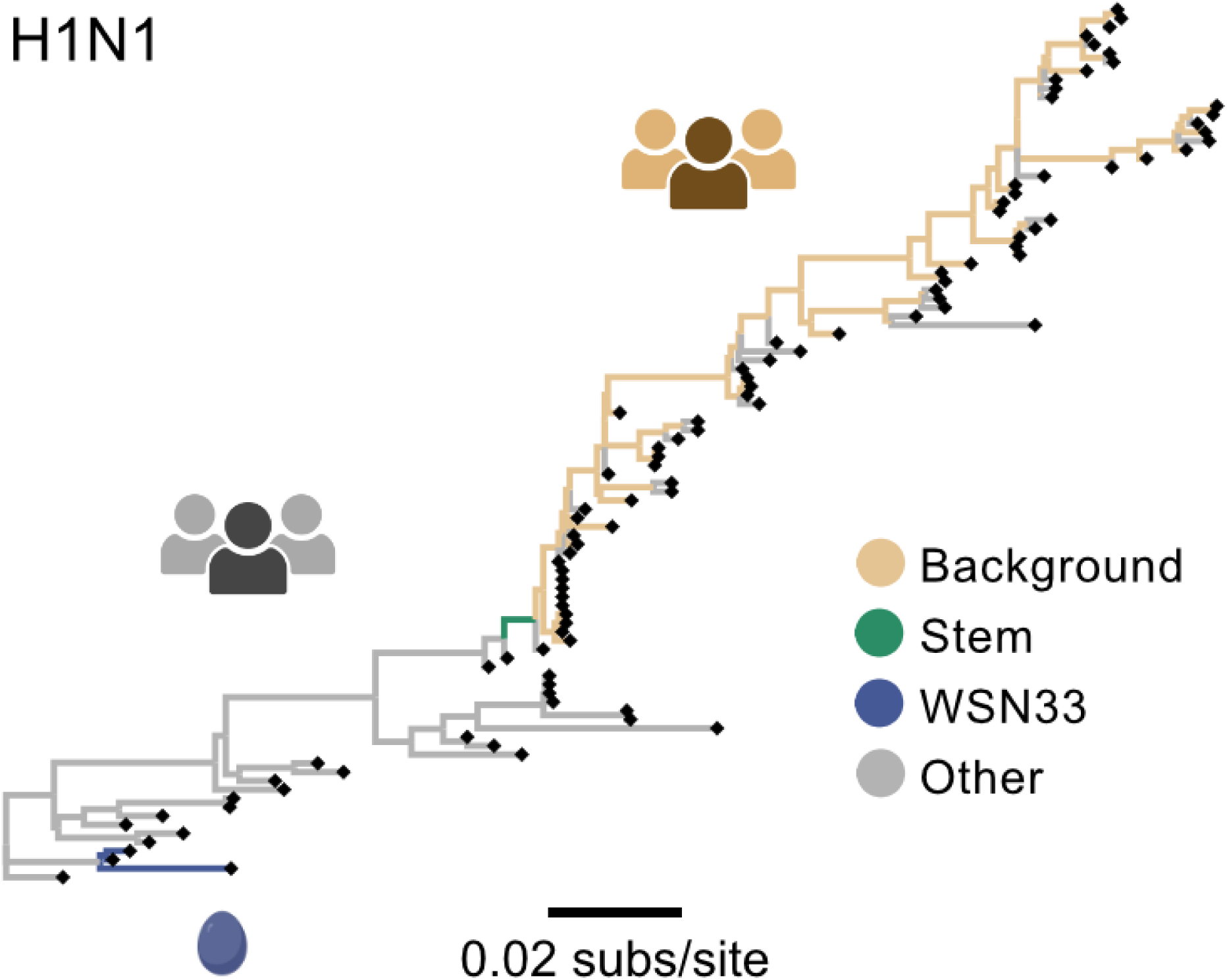
Phylogenetic tree of H1N1 HA segment, branch color indicates partition. The stem branch connects 1977 pandemic reemergence and basal ancestors, and is foreground for 1977 flu test. WSN33 (blue) branches are clade associated with laboratory passaged positive control. Post 1977 branches associated human to human transmission (brown) used as background. Other branches that are pre-1977 or tips that have a history of passaging (grey) are excluded from analysis. Icons created with BioRender.

**Supp Fig 3.**
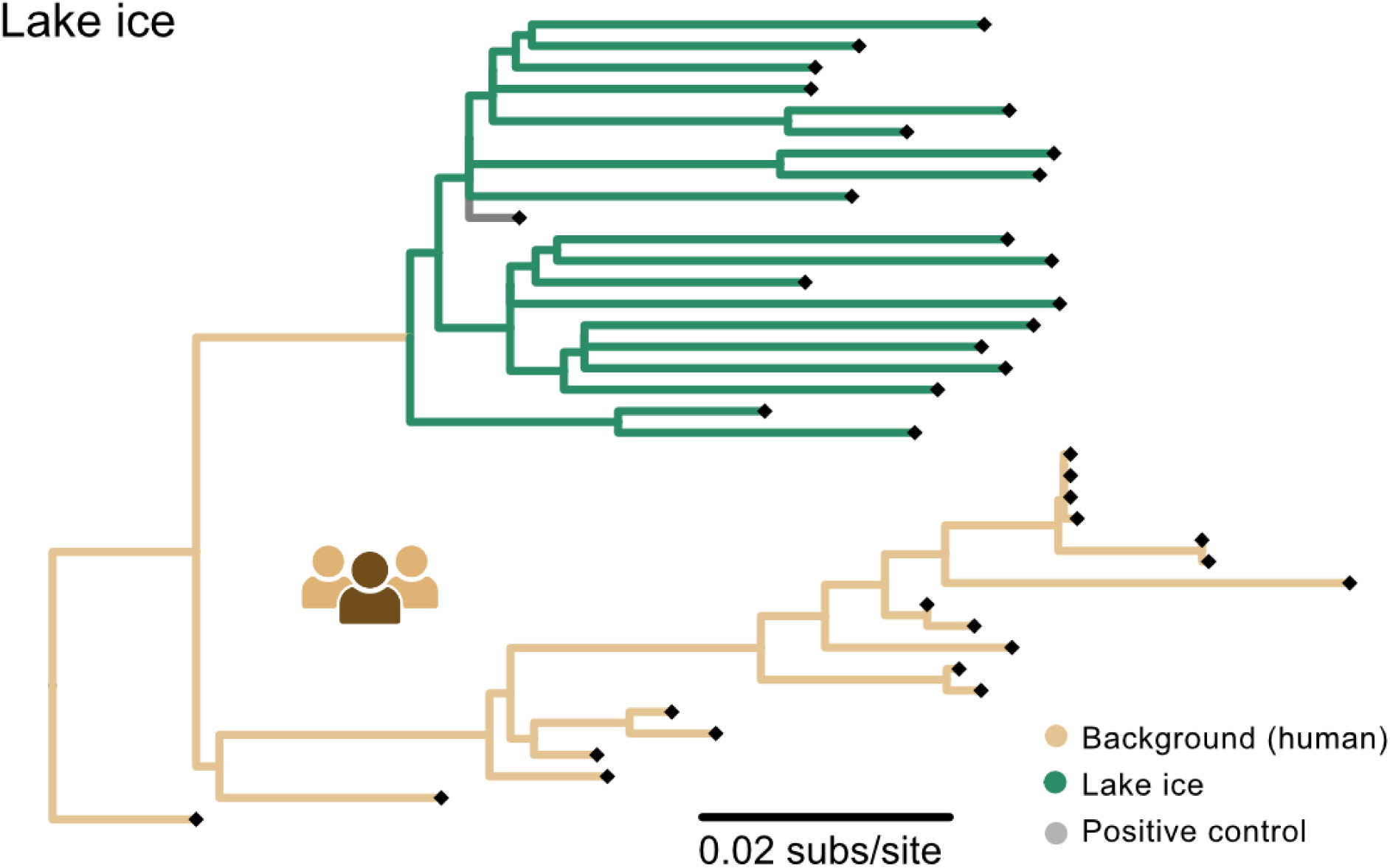
Phylogenetic tree of H1N1 HA segment, branch color indicates partition. Lake ice clade (green) is the foreground and branches associated with human to human transmission (brown) are background of Lake ice test. The study positive control (grey) is excluded. Icons created with BioRender.

**Supp Fig 4.**
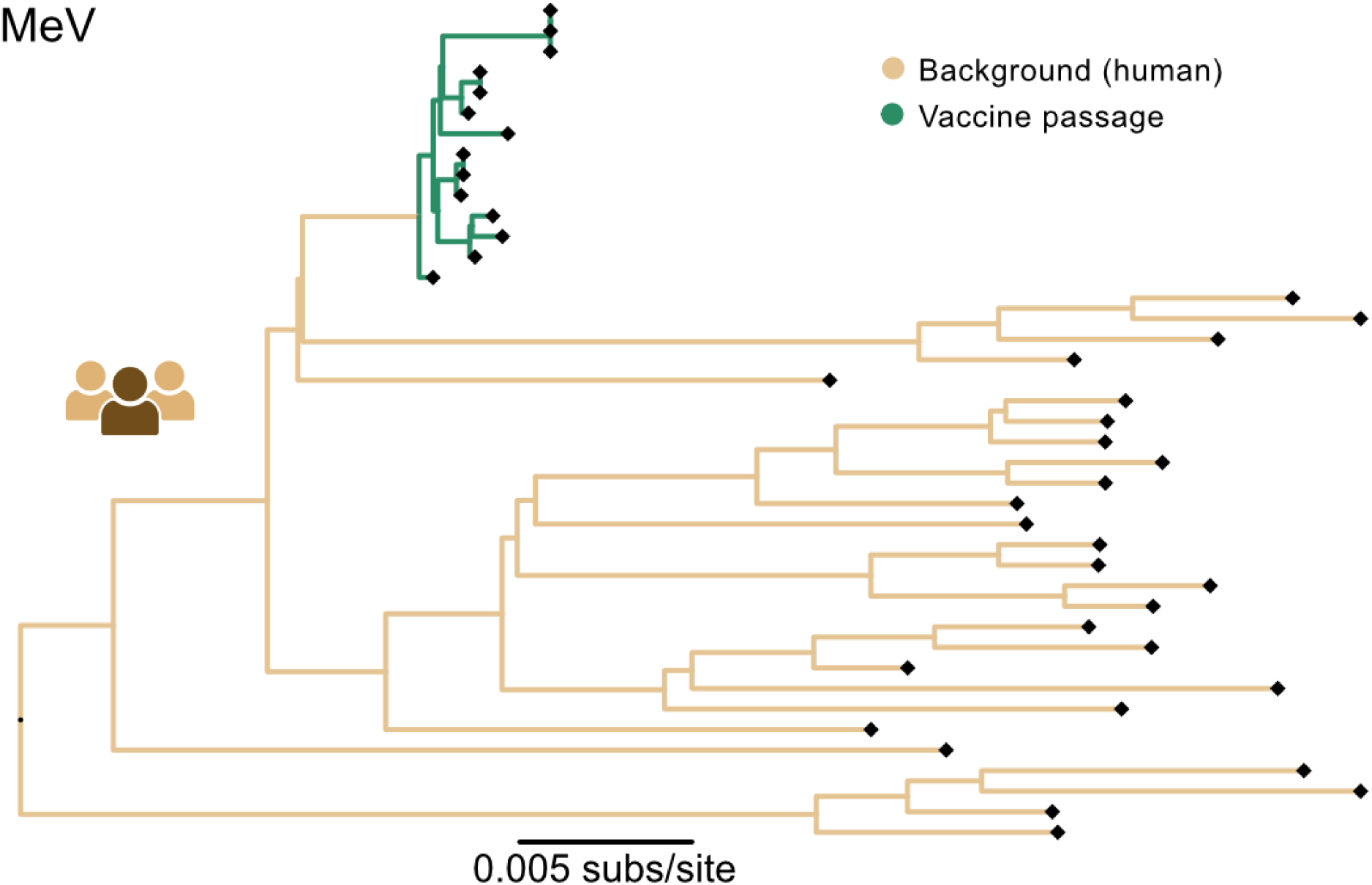
Phylogenetic tree of MeV genome, branch color indicates partition. Vaccine attenuated by passage (green) is the foreground and branches associated with human to human transmission. Icons created with BioRender.

**Supp Fig 5.**
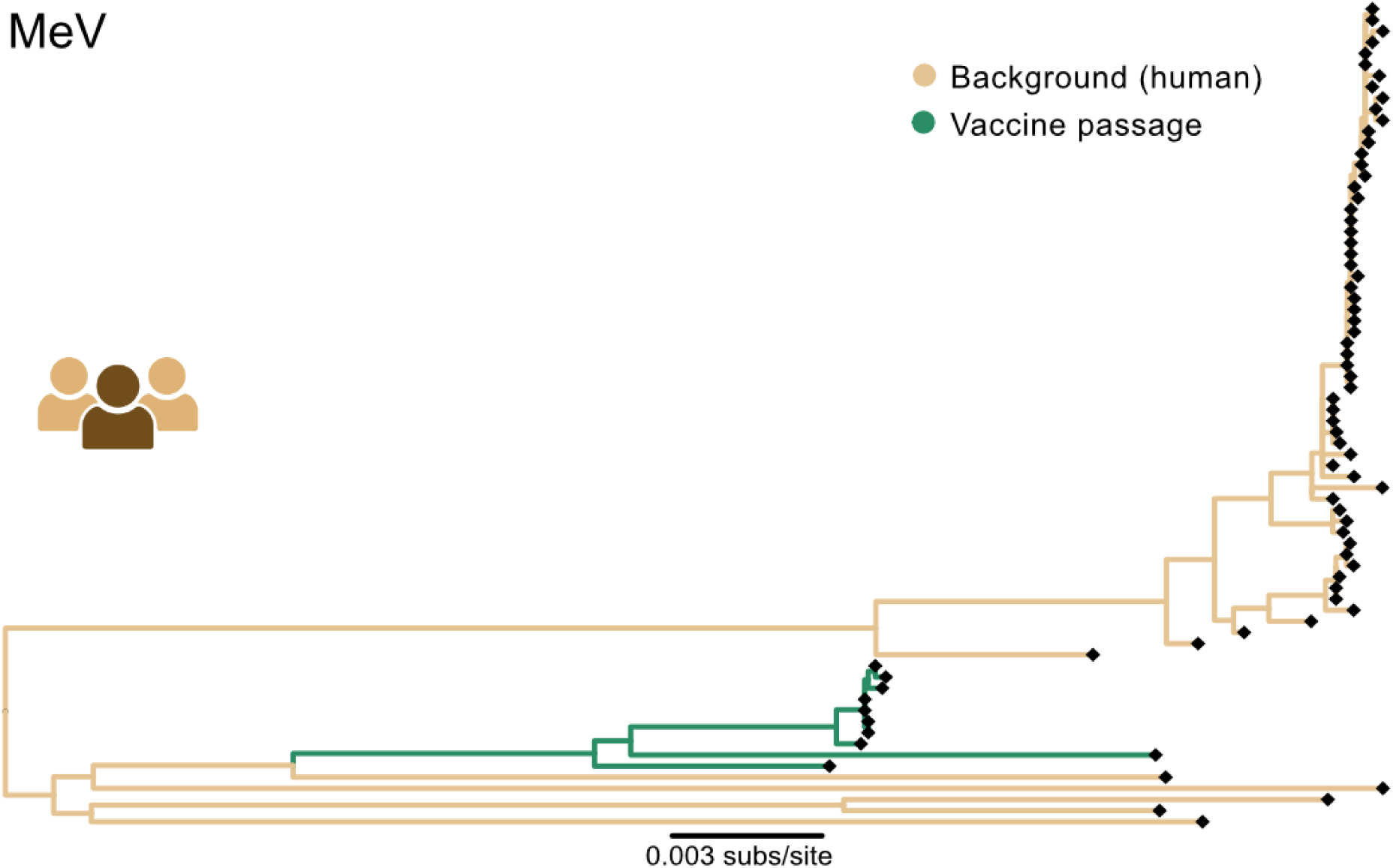
Phylogenetic tree of MuV genome, branch color indicates partition. Vaccine attenuated by passage (green) is the foreground and branches associated with human to human transmission. Icons created with BioRender.

**Supp Fig 6.**
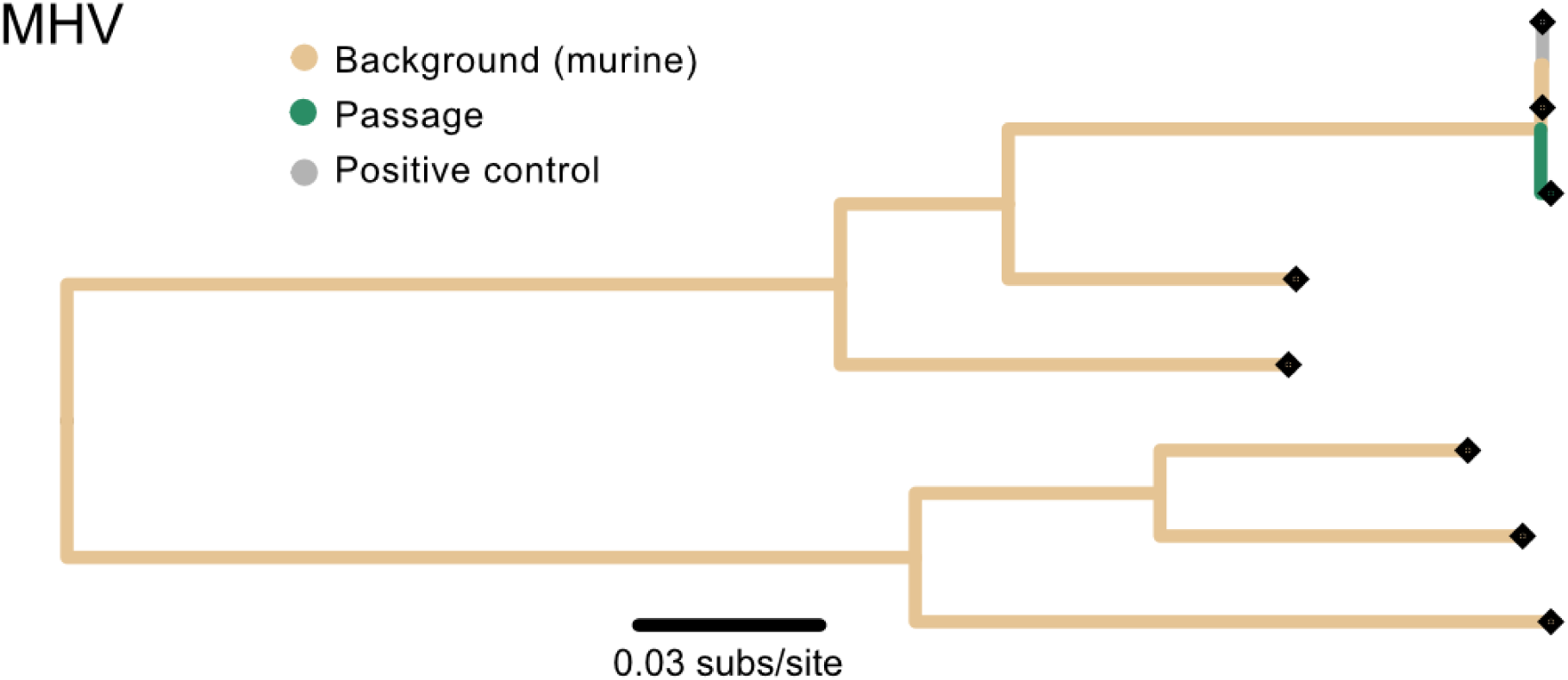
Phylogenetic tree of MHV non-recombinant region 07, defined by GARD, branch color indicates partition. Laboratory passaged virus (green) is the foreground and branches between murine strains (brown) are background of Lake ice test. The isolate used at the start of passaging (grey) is excluded.

**Supp Fig 7.**
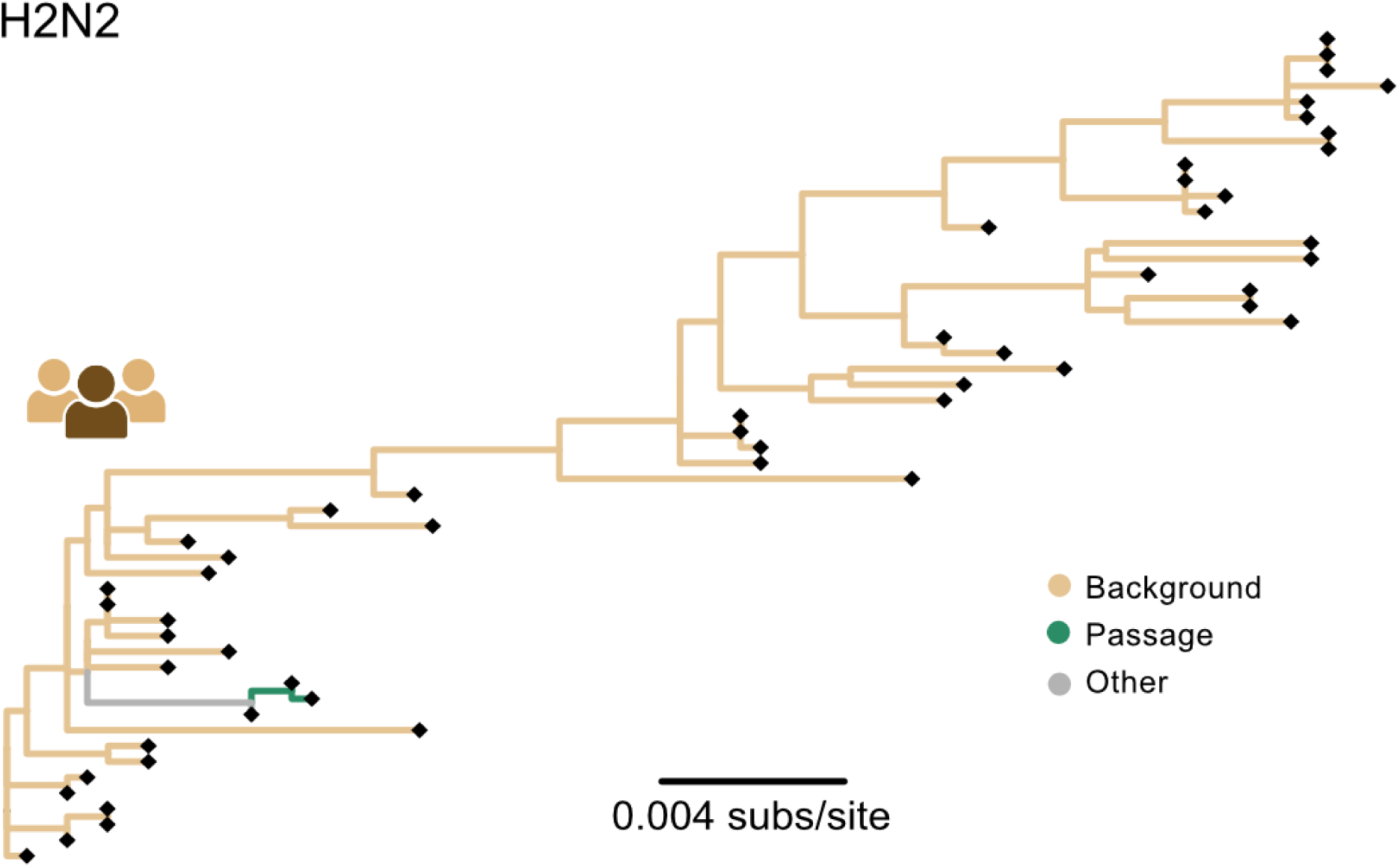
Phylogenetic tree of H2N2 PB1 segment, branch color indicates partition. Cold adaptation passage (green) is the foreground and branches associated with human to human transmission (brown) are background. The stem to isolate used at the start of passaging (grey) is excluded. Icons created with BioRender.

**Supp Fig 8.**
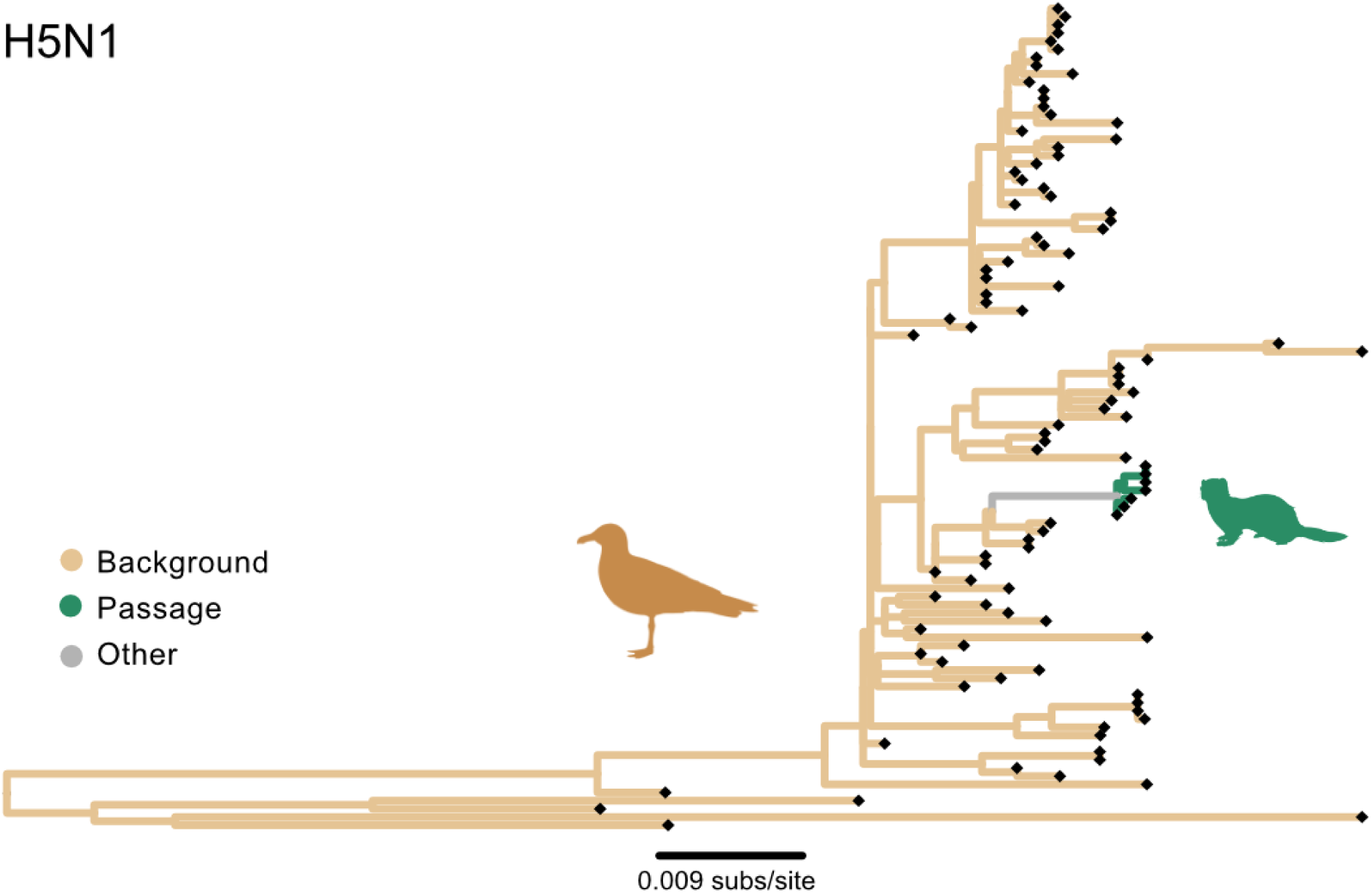
Phylogenetic tree of H5N1 PB1 segment, branch color indicates partition. Airborne ferret adaptation passage (green) is the foreground and branches associated with avian host reservoir (brown) are background. The stem to isolate used at the start of passaging (grey) is excluded. Icons created with BioRender.

